# Prevalence and determinants of neonatal danger signs in northwest Ethiopia: a multilevel analysis

**DOI:** 10.1101/597245

**Authors:** Tariku Nigatu, Abebaw Gebeyehu, Alemayehu Worku, Gashaw Andargie, Zemene Tigabu

## Abstract

**Background:** There is association between neonatal danger signs and neonatal deaths. Hence, understanding the factors associated with the occurrence of neonatal danger signs help reduce the stagnating neonatal mortality in countries like Ethiopia.

**Method:** A cross sectional community and facility linked study was conducted in 39 kebeles in Amhara region, North Gondar Zone of Ethiopia from March 3-18, 2016. A representative sample of 1,150 mother-newborn pairs were included in the study. Percentage was used to calculate the prevalence. Multilevel analysis was used to identify individual and kebele level characteristics associated with the occurrence of neonatal danger signs.

**Result:** The result showed that around a quarter, 286 (24.9%), of the newborns experienced one or more danger signs during the neonatal period. Significant differences were found between groups/kebeles in the occurrence of danger signs. At individual level, having low birth weight (AOR= 0.65; 95% CI: 0.48-0.88) and maternal danger signs during pregnancy and delivery (AOR= 1.93; 95% CI: 1.41-2.65) were found to be significantly associated with the occurrence of neonatal danger signs. At group/kebele level, antenatal care coverage (AOR= 0.35; 95% CI: 0.13-0.93) and year of health extension workers experience (AOR= 0.91; 95 % CI: 0.84-0.99) were significantly associated with the occurrence of neonatal danger signs.

**Conclusion:** The prevalence of neonatal danger signs is high. There are **i**ndividual and kebele level characteristics associated with occurrence of danger signs in newborns. Expanding maternal health services and strengthening the health extension program is critical.

## Background

The neonatal period marks the transition from intrauterine to extra uterine life. Usually the transition is smooth. Sometimes however, the process can be complicated leading to neonatal mortality(1).

Every day, an estimated 7,700 newborns die globally(2). The vast majority of these deaths happen in resource limited settings including Ethiopia(3). Linked with the high prevalence of home delivery in developing countries, most neonatal deaths occur at home(4).

Even though global efforts halved neonatal mortality from 4.7 million to 2.8 million between 1990 and 2013(5), the contribution of neonatal deaths to under five childhood mortality consistently grew from 37% to 44% in the same period(6)(7). The reason being a slow decline in neonatal mortality compared to under-five deaths(6)(8).

Similar trends are also observed in Ethiopia. According to the Ethiopian Demographic and Health Survey (EDHS), neonatal deaths declined from 49 deaths per 1000 live births in 2000 to 29 deaths per 1000 live births in 2016(9). However, the reduction was slow and sometimes stagnant resulting in a growing contribution to under-five mortality(10).

UNICEF and WHO identify the following nine symptoms as danger signs in newborns: 1) Not feeding since birth or stopped feeding, 2) Convulsions, 3) Respiratory rate of 60 or more, 4) Severe chest in-drawing, 5) Temperature ≥ 37.5° C, 6) Temperature ≤ 35.5° C, 7) movement only when stimulated or not even when stimulated, 8.)Yellow soles (sign of jaundice) and signs of local infection and, 9) Reddened or pus draining umbilicus, skin boils, or pus draining eyes(11).

These danger signs in newborns are nonspecific and each danger sign can be a sign of almost any disease or illness (12). Lack of knowledge about these neonatal danger signs is also a major barrier to treatment seeking (13–16), which may ultimately lead to neonatal death(12,17).

Better understanding of neonatal danger signs and the factors that affect their occurrence is very important to help reduce neonatal mortality. Evidence is also scarce on the factors responsible for newborn danger signs. This study is conducted to determine the prevalence of and identify factors operating at multiple levels; individual and contextual factors; that are associated with the occurrence of neonatal danger signs.

## Methods

### Study design, setting and source population

A cross sectional community and facility linked study was conducted from March 3-18, 2016 in North Gondar Zone of Ethiopia. North Gondar is in Amhara region located in the northwest part of Ethiopia (18). The zone has 24 woredas (districts). According to the Central Statistical Agency (CSA), in 2017, the total projected population in the zone based on the 2007 national population and housing census was 3,654,920 of which 1,847,631 (51%) were males (CSA, 2017). As of 2016, the zone has 9 government hospitals, 126 health centers and 563 health posts. There are also many private clinics most of them located in urban areas.

### Study population

Women who delivered live babies in the past six months. Health extension workers and health posts found in the study area were included in the study. Health extension workers (HEWs) are female, salaried, frontline health workers that provide basic health promotion and disease prevention services to rural communities in Ethiopia. The HEWs provide the services in house to house visit and at a health post.

### Variables

#### Outcome variable

The occurrence of neonatal danger signs during the first 28 days of life was the outcome variable.

#### Independent variables

Individual and kebele level characteristics were examined for possible associations with neonatal danger signs. Kebele is the smallest political administrative unit below district with an estimated average population size of 5000. The conceptual framework depicting the assumed relationships between kebele and individual level characteristics with the outcome variable is shown below (Figure 1).

**Figure 1:**
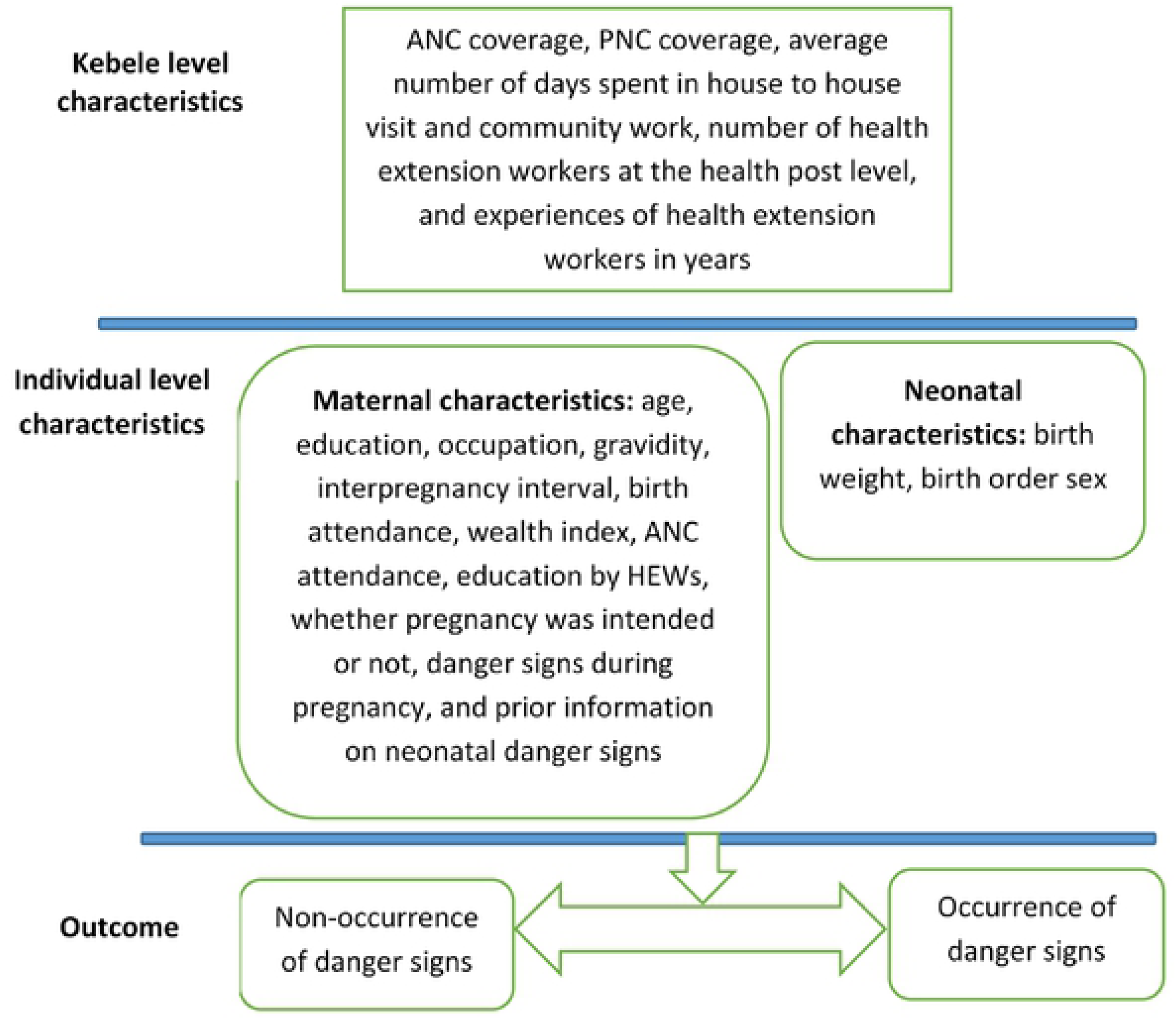
Conceptual framework for individual & Kebele-level determinants influencing neonatal danger signs

#### Individual level variables

Individual-level variables included maternal and neonatal characteristics. Maternal characteristics included: maternal age, occupation, exposure to health education, gravidity, insecticide treated nets (ITN) use during the last pregnancy, whether the pregnancy of the indexed child was wanted or not, occurrence of maternal danger signs during pregnancy and the uptake of skilled birth attendance during the last delivery. Newborn characteristics were measured based on the reports of the mother. These included; occurrence of danger signs in the newborn, birth weight, and sex of the newborn.

In settings where children are not often weighed at birth, the mothers’ report of the size of their babies at birth is used as the proxy for the child’s weight(10). In this study, birth weight of the newborns was measured by asking mothers to rate the birth weight of the indexed newborn as very small, small, normal, big and very big. Newborns rated as very small and small were re-categorized as small and those rated normal and above were rated as normal during analysis. In addition, household wealth index was constructed using principal components analysis. The wealth index was weighted for urban and rural areas before producing the combined wealth index.

#### Kebele level variables

Groups of mothers and newborns are clustered within kebeles. There is one health post in each Kebele providing basic disease prevention and health promotion services for the kebele population. Health post level service coverage and other health related data were taken as kebele level characteristics. These included coverage of at least one antenatal and postnatal care, number of health extension workers (HEWs) working at each health post and the average number of days HEWs spend for house to house visit per week and the experiences of HEWs in years.

#### Sample size and sampling

Two population proportion formula was used to calculate sample size using statcalc in Epi-Info version 7.1.5.0. Variables taken from two studies were used to calculate the sample size(15,19). Sample sizes were calculated independently based on the two studies by assuming 95% confidence level (1-α), 80% power (1-β) and unexposed to exposed ratio of 1:1. The larger calculated sample size based was 388. With a design effect of 2, and non-response rate of 10%, the final sample size was 854 mother-newborn pairs. This study was part of a bigger study with a sample size of 2,158 mothers. Of this sample, 1,150 of the mothers had delivered a live baby in the past six months. Hence, the sample size was taken as sufficient for this study.

A multistage stratified cluster sampling technique was used to select the mothers. First, 39 kebeles were randomly selected from three districts (Debark, Dabat and Wogera) proportional to the size of kebeles in the woredas. Then, villages or ‘’Gotes’’ from the selected kebeles were randomly selected. Data was collected from mothers in all eligible households in the clusters or‘’Gotes’’. Kebele level data was collected from health posts service statistics reports and interviews with HEWs.

#### Data Collection

Fifty four data collectors and five supervisors, all of them with at least first degree in health, were recruited and received two day training. A pretested structured interviewer administered questionnaire was used to collect data on individual level characteristics. The questions were adapted from different surveys. Structured questionnaire was used for data collection from HEWs and health posts.

#### Data Processing and Analysis

The data was entered and cleaned using Epi-Info version 7.1.5.0. Data cleaning was made by running frequencies and descriptive statistics. The cleaned data was exported to STATA version 13 for analysis.

#### Descriptive analyses

First, exploratory descriptive statistics was applied to understand the nature of the data. The frequencies of each of the categories within the explanatory variables were calculated. The overall study area prevalence was also estimated. Multiple response analysis was conducted to determine the prevalence of each danger signs.

#### Modelling approaches

Two-level multivariable logistic regression was applied to account for the hierarchical nature of the data and to get unbiased estimate of regression coefficients. Three models were fitted in the analysis. Model one, the empty or unconditional model, decomposed the total variance in the dependent variable in to individual and group/kebele level variances. The portion of the total variance explained by kebele level characteristics was measured by the intra-class correlation coefficient (ICC). The chi-squared test that compares the empty nested model with the classical logistic model was used to test the significance of kebele level characteristics in explaining the total variance of the outcome variable. Model two included all the individual-level variables (maternal and neonatal). In model two, we assessed the compositional effect of kebeles in explaining the variance among the groups (kebeles). Model three encompassed the combined effect of individual and kebele-level characteristics. With the third model, we determined the significance of individual and kebele level characteristics in explaining the variances among the groups.

#### Fixed effects

The relationships between individual and kebele-level characteristics with the occurrence of neonatal danger signs were reported in term of odds ratios with p-values at 95% confidence interval.

#### Random effects

Random effects that measure the variations among the groups/kebeles were expressed in terms of Intra-class correlation (ICC).

#### Model fitness & precision

The log likelihood of the models were estimated to assess the fitness of the model relative to the other models. Variance Inflation Factor was used to test the presence of multicollinearity in the model. Stata software package of version 14 was used for the analyses. Statistical significance of the predictor variables were determined by two tailed Wald test at a 5% level of significance.

#### Ethical Considerations

The study was reviewed and approved by the University of Gondar Institutional Review Board (IRB). Permission was obtained from all kebeles in the study areas. During data collection, study subjects were asked for oral informed consent. For adolescent mothers below the age of 18, informed consent was taken as per the National Research Ethics Review Guideline’s recommendation for emancipated minors and with the approval of the IRB. All study participants were invited to participate voluntarily in the study. In addition, they were informed on the potential benefits, harms, the confidentiality and the possibility of withdrawing from the interview even without giving reasons. All interviews were conducted in private settings.

## Result

### Socio-demographic characteristics

A total of 1,150 mother-newborn pairs were included in the study. Most, 1,059(92.1%), of the mothers resided in rural areas. The mean (±SD) and median (IQR) age of the mothers were 27.5(±6.7) and 27(10) years, respectively.

Nearly a third, 393 (34.1%), of the mothers were young aged 15-24 years. Farming was the main, 1,066(92.7%), source of livelihood in the study areas. The majority, 743(64.6%) and 938 (81.6%), of the mothers were illiterate and had history of two or more pregnancies, respectively. Most of the deliveries, 713(62.0), for the indexed newborns were attended by unskilled birth attendants. Nearly all of the pregnancies for the indexed newborns, 1,126(98%), were spaced less than 24 months. More than a quarter, 271(23.6%), of these pregnancies were unintended (Table 1).

**Table 1:** General characteristics of the study population: individual characteristics; northwest Ethiopia, March 2016

| Variables | Number (%) | Danger sign |  |
| --- | --- | --- | --- |
|  |  | Yes | No |
| **Maternal characteristics** |  |  |  |
| **Wealth index** |  |  |  |
| Poorest | 217 (18.9) | 72 | 145 |
| Poor | 230(20.0) | 55 | 175 |
| Medium | 234(20.3) | 60 | 174 |
| Rich | 221(19.2) | 41 | 180 |
| Richest | 248(21.6) | 58 | 190 |
| **Age** |  |  |  |
| 15-19 | 119(10.3) | 33 | 86 |
| 20-24 | 274(23.8) | 62 | 212 |
| 25-29 | 292(25.4) | 77 | 215 |
| >30 | 465(40.4) | 114 | 351 |
| **Educational status of mother** |  |  |  |
| Illiterate | 743(64.6) | 188 | 555 |
| Able to read and write | 38(3.3) | 13 | 25 |
| 1-4<sup>th</sup> grade | 117(10.2) | 32 | 85 |
| 5-8<sup>th</sup> grade | 149(13.0) | 29 | 120 |
| 9-10<sup>th</sup> grade | 84(7.3) | 18 | 66 |
| 11-12<sup>th</sup> grade | 8(0.7) | 2 | 6 |
| Higher education | 11(1.0) | 4 | 7 |
| **Mothers Occupation** |  |  |  |
| Farmer | 1066(92.7) | 263 | 803 |
| Government employee | 12(1.0) | 4 | 8 |
| Other (merchant, daily laborer) | 72(6.3) | 19 | 53 |
| **Gravidity** |  |  |  |
| 1 | 212(18.4) | 55 | 157 |
| 2-5 | 709(61.7) | 170 | 539 |
| >6 | 229(19.9) | 61 | 168 |
| **Birth Attendance** |  |  |  |
| Unskilled (including HEWs) | 713(62.0) | 187 | 526 |
| Skilled | 437(38.0) | 99 | 338 |
| **Pregnancy was unintended** |  |  |  |
| Yes | 879(76.4) | 210 | 669 |
| No | 271(23.6) | 76 | 195 |
| **Health education by health extension workers** |  |  |  |
| Yes | 518(45.0) | 108 | 410 |
| No | 632(55.0) | 178 | 454 |
| **Heard about danger signs before** |  |  |  |
| Yes | 474(41.2) | 92 | 382 |
| No | 676(58.8) | 194 | 482 |
| **Neonatal characteristics** |  |  |  |
| **Sex of newborn** |  |  |  |
| Female | 571(49.7) | 139 | 432 |
| Male | 579(50.3) | 147 | 432 |

| **Birth weight** |  |  |  |
| --- | --- | --- | --- |
| Small | 611 (0.53) | 174 | 437 |
| Normal | 539 (0.47) | 112 | 427 |
| **Birth order** |  |  |  |
| 1<sup>st</sup> | 215 (0.19) | 53 | 162 |
| 2<sup>nd</sup> | 194 (0.17) | 53 | 141 |
| 3<sup>rd</sup> | 182 (0.16) | 41 | 141 |
| 4<sup>th</sup> | 183(0.16) | 39 | 144 |
| ≥ 5<sup>th</sup> | 376(0.33) | 100 | 276 |

The percentage of male and female newborns was almost the same. A little more than half, 611 (53%), of the newborns were rated to have small birth weight at the time of birth. A third of, 376(33%), the newborns were either the 5^th^ or more children in the family (Table 1).

The coverages of maternal health services (antenatal care (ANC), skilled delivery and postnatal care), and the average number of health extension workers, their experience in years and the average number of days they spent in house to house visit and community level health related activities is shown below (Table 2).

**Table 2:** General characteristics of the study population: Kebele level variables

| Health Post level characteristics | Mean (± SD) |
| --- | --- |
| At least one or more ANC coverage in the Kebele | 0.65 (±.23) |
| PNC coverage in the Kebele | 0.59 (±.23) |
| Skilled birth attendance | 0.43(±.22) |
| Average number of days HEWs spend in the community | 2.3 (±1.46) |
| HEWs average year of experience in years | 7.78(±2.89) |
| Number of HEWs working in the Kebele | 2.05(±.55) |

### Prevalence of neonatal danger signs

The mothers reported that around a quarter, 286 (24.9%), of the newborns experienced one or more of the WHO defined danger signs during the neonatal period. Mothers from the poorest households reported the highest percent of cases of neonatal dangers signs (Table 1).

The commonest neonatal danger sign reported was fever, 213(74.5%), followed by fast breathing and difficulty breathing, 127 (44.4%) and 107 (37.4%), respectively. The least reported danger sign was yellow soles and feet (jaundice), 14 (4.9%) (Table 3).

**Table 3:** Distribution of neonatal danger signs, northwest Ethiopia, March 2016

| Danger signs | Responses |  | Percent of Cases |
| --- | --- | --- | --- |
|  | N | Percent |  |
| Fever | 213 | 28.00 | 74.50 |
| Hypothermia | 48 | 6.30 | 16.80 |
| Fast breathing | 127 | 16.70 | 44.40 |
| Difficulty of breathing | 107 | 14.10 | 37.40 |
| Red swollen and pusy eye | 55 | 7.20 | 19.20 |
| Red swollen and pusy umbilicus | 65 | 8.50 | 22.70 |
| Convulsion | 22 | 2.90 | 7.70 |
| Floppiness and absence of movement | 28 | 3.70 | 9.80 |
| Yellow soles and feet (jaundice) | 14 | 1.80 | 4.90 |
| Inability to suck | 82 | 10.80 | 28.70 |
|  | 761 | 100.00 | 266.10 |

### Random effects

Significant heterogeneity was observed among kebeles. The ICC calculated based on the null or empty model was significant at 0.094 implying that 9.4% of the total variance in the occurrence of neonatal danger signs was attributed to the differences among kebeles/groups. This also implies that the correlation between newborns living in the same kebele in the likelihood of having neonatal danger sign was 0.09.

A significant reduction in kebele-level variance was observed in model two. This indicated the significance of the compositional effect (individuals within the kebeles) in explaining the between group variance.

However, we extended model two by introducing kebele level characteristics to form model three. In the final model (model three), kebele-level variance was significantly reduced further after adjusting for both individual and community-level characteristics.

### Fixed effects

Fixed effects of model two show the associations between individual-level characteristics and the occurrence of neonatal danger signs when kebele-level characteristics were not considered. Fixed effects of model three show the associations of both individual and kebele-level characteristics with neonatal danger signs.

After considering both individual and kebele-level characteristics in model three, it was observed that newborns with normal birth weight were 35 percent less likely to experience neonatal danger signs (AOR 0.65; 95% CI 0.48-0.88) compared to small birth weight newborns. Other newborn level characteristics in the study were not significantly associated with neonatal danger signs.

Similarly, a maternal level characteristic was also found to be significantly associated with neonatal danger signs. Newborns delivered by mothers who experienced one or more danger signs during pregnancy and delivery had 93% higher odds of having neonatal danger signs compared to newborns delivered from mothers who did not experience danger signs themselves (AOR 1.93; 95% CI 1.41-2.65).

Some kebele level characteristics were also significantly associated with the occurrence of neonatal danger signs in newborns. Antenatal care coverage of the kebele and year of health extension workers experience were associated with neonatal danger signs (AOR 0.35; 95% CI 0.13-0.93 and AOR 0.91; 95 % CI 0.84-0.99), respectively (Table 4).

**Table 4:**
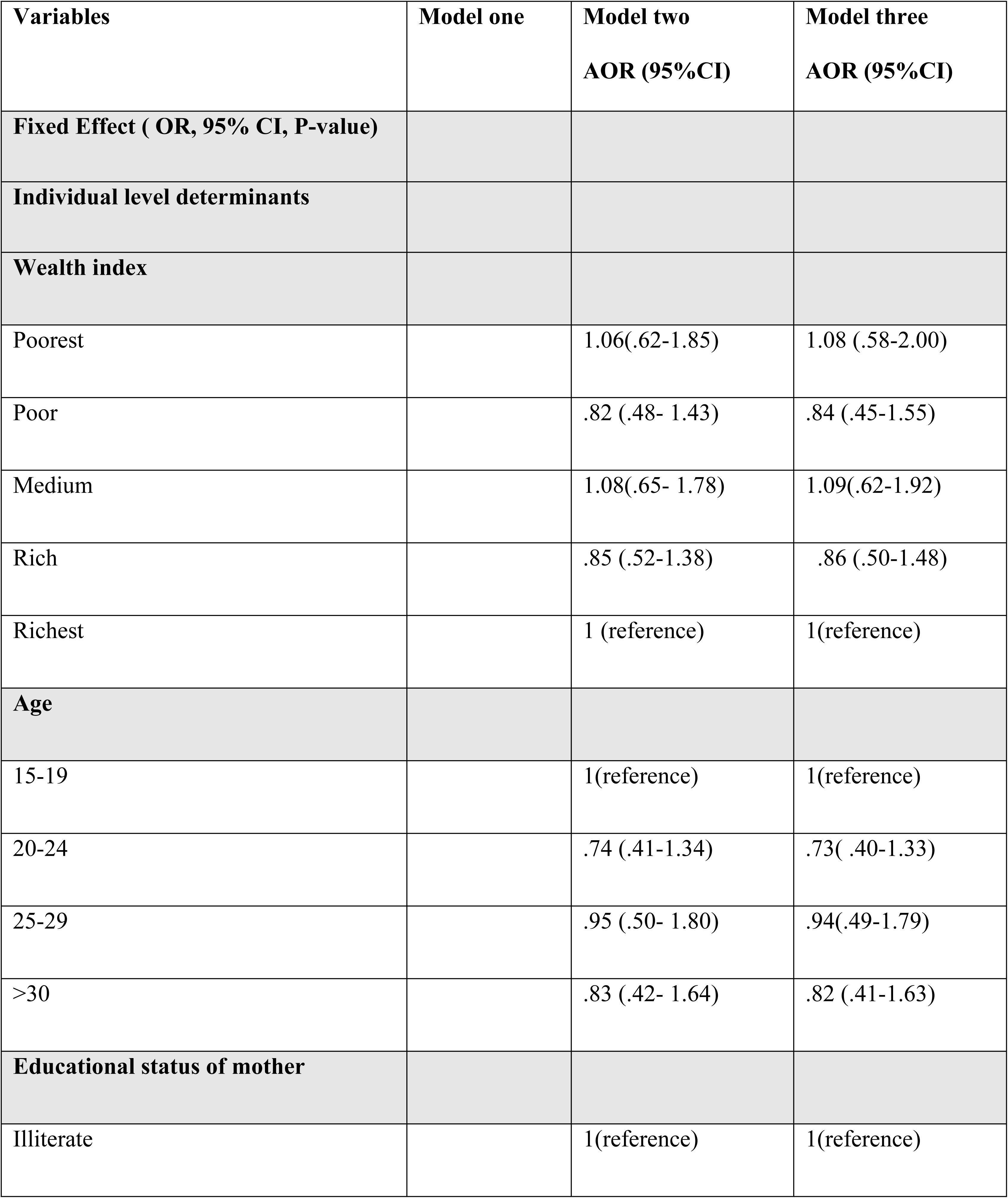
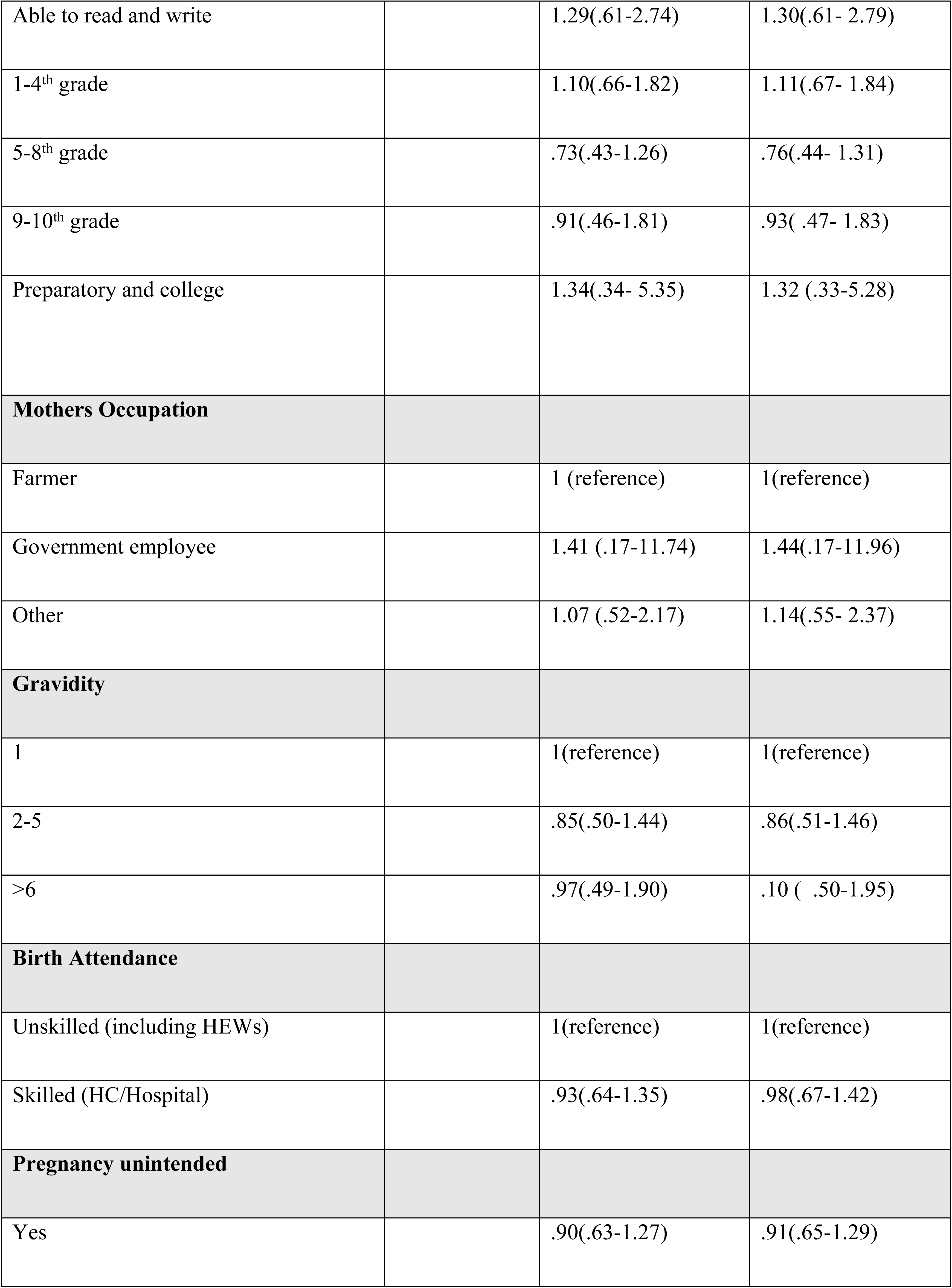
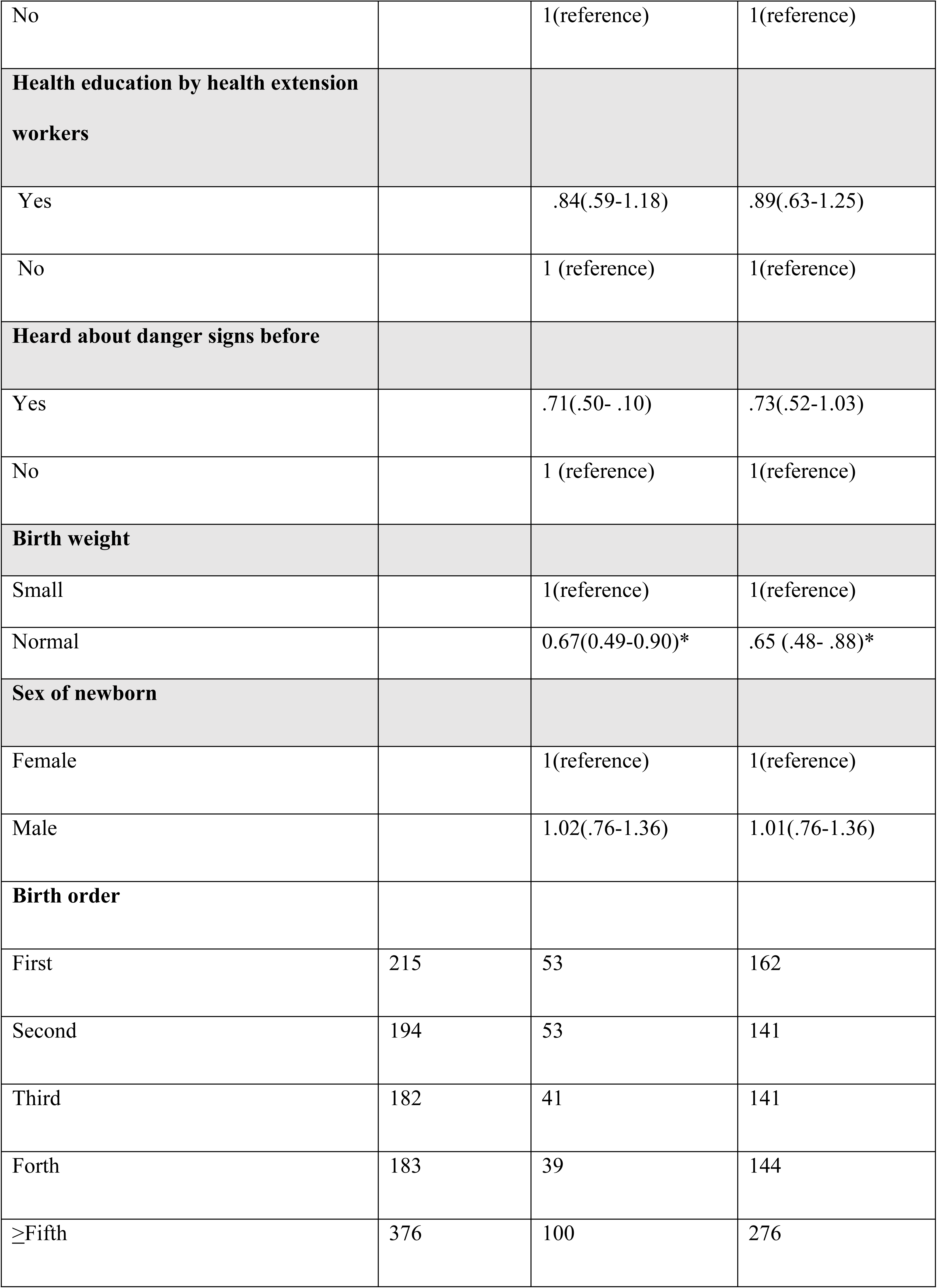
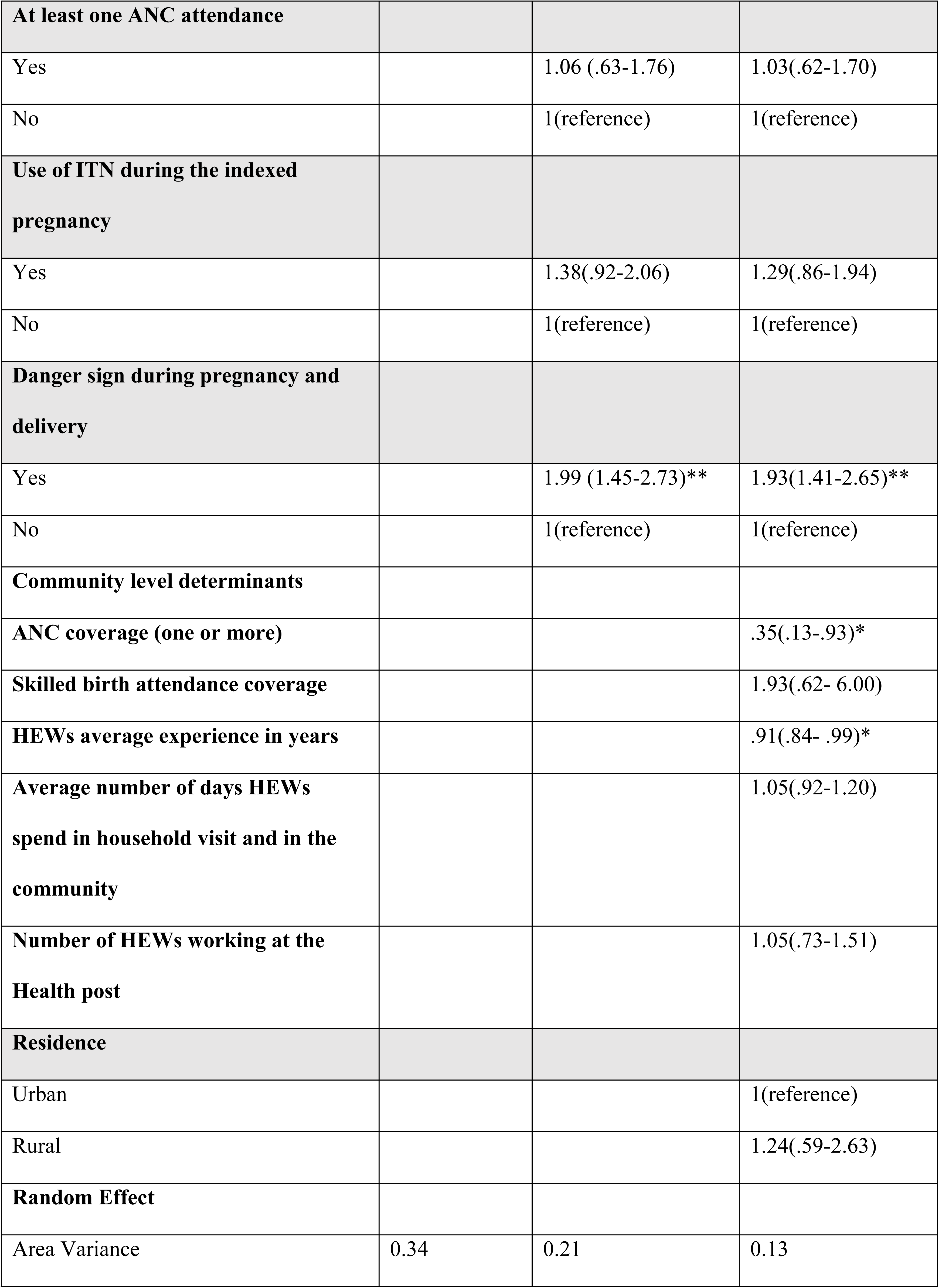
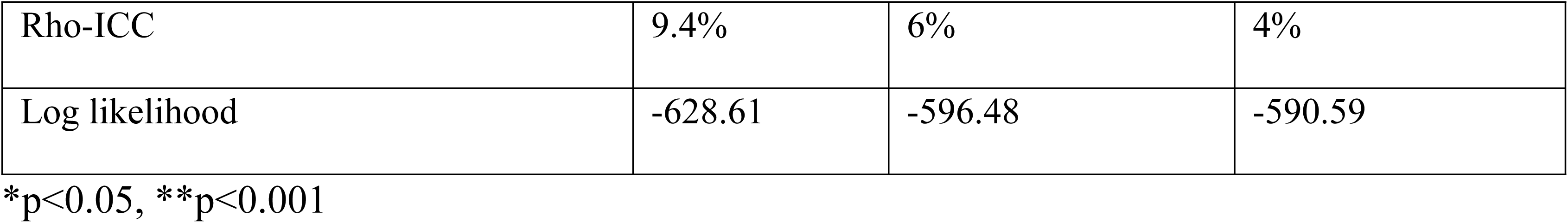
Associations between neonatal mortality and individual and community level determinants

### Model fit statistics

There was a progressive increase in the negative log likelihood observed in model one, model two and model three. This implies that model three explained the determinants better than either model one or two.

## Discussion

This study investigated the association of maternal, neonatal and kebele level characteristics with the of occurrence of neonatal danger signs. It also tried to determine the prevalence of danger signs in general and specific danger signs in particular among newborns in the study area.

In this study, more than 90 percent of the variability in the occurrence of danger signs in newborns was explained by individual characteristics. This shows that individual level characteristics (the kebele composition) were more important than the group/kebele level characteristics in determining the occurrence of danger signs in newborns.

The prevalence of neonatal danger signs was found to be 25%, which means that one in every four newborns in the study area experienced one or more danger signs in the first 28 days of life. This implies a significant burden of morbidity among the most vulnerable member of human beings; newborns. Similar finding was also reported in a study conducted in Ghana(20).

Both individual and group level characteristics were significantly associated with the occurrence of the danger signs. At individual level, birth weight of the newborn, as judged by the mother, was found to be an important factor in predicting the occurrence of neonatal danger signs. Even if there were no prior studies found on predictors of neonatal danger signs in particular, many studies showed that low birth weight is an important predictor of neonatal mortality (17,21–23).

Newborns from mothers that experienced danger signs during the indexed pregnancy and delivery were associated with higher odds of having danger signs in the newborns. This indicates that danger signs during pregnancy are important predictors of danger signs in newborns. This strengthens the need to follow up mothers with danger signs to avoid undesired outcome of both the mother and the newborn.

Coverage of at least one antenatal care was significantly associated with reductions in neonatal danger signs. Studies in Ethiopia and Kenya showed that mothers that attended antenatal care were more knowledgeable about neonatal danger signs than mothers that did not(24,25). Mothers with better knowledge of neonatal danger signs also tend to have better health and care seeking behavior (14). Hence, the effect of antenatal care attendance on the occurrence of neonatal danger signs in this study could be because of better knowledge of danger signs among mothers that attended antenatal care, which might have affected better care seeking behavior resulting in reduction in the occurrence of neonatal danger signs.

Health extension workers experience was negatively associated with the occurrence of danger signs in newborns. Long years of experience could mean better knowledge of the area, culture and care seeking behavior of the community. This knowledge of the area might have helped the health extension workers to plan and implement health promotion and disease prevention activities that positively impact the occurrence of danger signs in newborns.

## Conclusion

This study demonstrated that the burden of illness among newborns is high in the study area. It also revealed that both individual and kebele level characteristics determine the occurrence and non-occurrence of danger signs in newborns. Individual level characteristics (the kebele composition) were also found to be more important than the group/kebele level characteristics in determining the occurrence of danger signs in newborns.

Improving coverage of maternal health services, particularly antenatal, delivery and postnatal care, is important to reduce neonatal danger signs and thereby reduce the associated neonatal mortality. Educating mothers about neonatal danger signs during pregnancy and delivery and strengthening the health extension program is critical. Most importantly, strengthening postnatal home visits of both the mother and the newborn is important to identify and treat newborn danger signs early.

### Limitation of the study

One of the limitation of this study could be a recall bias associated with the length of time mothers were expected to report their experience. To reduce this bias, the interviewers mentioned each danger signs one by one and gave mothers adequate time to respond. A prospective cohort study may give more estimates of the determinant of newborn danger signs.

### List of Abbreviations

ANC: Antenatal Care
AOR: Adjusted Odds Ratio
CBNC: Community Based Newborn Care
CI: Confidence Interval
EDHS: Ethiopian Demographic and Health Survey
HEP: Health Extension Program
HEWs: Health Extension Workers
ICC: Intra-Class Correlation Coefficient
iCCM: integrated Community Case Management
IQR: Interquartile Range
ITN: Insecticide Treated Nets
SD: Standard Deviation
SDG: Sustainable Development Goal

## Declarations

### Ethics approval and consent to participate

This study received ethical clearance from the University of Gondar Institutional Review Board (IRB), Ethiopia. Permission was obtained from kebele administrations. Verbal informed consent was obtained from the study participants. For adolescent mothers below the age of 18, informed consent was also taken as per the National Research Ethics Review Guideline’s recommendation for emancipated minors and with the approval of the IRB. This method of data collection was approved by the IRB of the University of Gondar

### Consent for publication

Not applicable

### Availability of data and Materials

The dataset contains individuals’ private information and can’t be shared publicly. However, data can be made available from the corresponding author and up on permission of the University of Gondar based on reasonable requests.

## Authors’ contributions

TN conceived and designed the study, collected data, performed the statistical analysis and drafted the manuscript. AG helped in the conceptualization, design, coordination, and revision of the manuscript. GA designed and coordinated the study, and revised the manuscript. ZT coordinated the study and revised the manuscript. AW helped in the design, analysis and revision of the manuscript. All authors read and approved the final manuscript.

## Acknowledgement

the authors of this paper would like to thank the research participants, Dabat Demographic Health Surveillance site (DHSS) and the local administrations in the study areas for their support during the conduct of this study.

